# MEAM-OWL for browser-based analysis of motif enrichment and associations with gene expression

**DOI:** 10.64898/2026.11.17.767221

**Authors:** Xin Jin, Kota Kambara, Sintho Wahyuning Ardie, Daisuke Tsugama

## Abstract

**Motivation:** Identifying functional regulatory motifs (such as *cis*-regulatory elements) remains challenging because conventional motif enrichment relies on binary comparisons between target and background sequences, overlooking quantitative relationships between motif copy number and gene expression or other quantitative traits. Existing tools for such analysis require specific and inconsistent input formats, incompatible with a streamlined workflow.

**Results:** We present MEAM-OWL (Motif Enrichment Analysis Modules on the Web, Lite) to overcome those issues. As well as standard target-versus-background enrichment, MEAM-OWL can evaluate motif enrichment on the basis of either expression-ranked lists or Pearson correlation tests between motif counts and expression levels. These analyses can be preceded by sequence extraction and ingestion from genome and genome annotation files, and can be followed by annotation of detected motifs with known *cis*-elements from public databases. In pearl millet under dehydration, consistent with previous findings, MEAM-OWL detected the positive correlation between gene induction levels and the copy numbers of the drought-responsive ABRE (abscisic acid response element)-related motifs in their promoters. Furthermore, MEAM-OWL uncovered the correlations between the copy numbers of the VIP1-binding element in the promoters of *Arabidopsis thaliana* and the expression changes of mechanical stress-responsive genes across lines with perturbed expression of the transcription factor VIP1 and its homologs, as well as its enrichment that could be detected by a conventional tool. These results demonstrate that integrating quantitative expression-motif correlations can promote discoveries of biologically meaningful regulatory relationships and seamlessly connect them to hypothesis building and wet-lab validation. MEAM-OWL can be used at https://tsugama-eng.github.io/meam-owl/.

## 1 Introduction

Short, conserved DNA and RNA sequences such as *cis*-regulatory elements can govern gene expression and other biological processes. To identify such functional sequences (or motifs), several computational tools have been widely adopted, including HOMER [1], STREME [2] (in the MEME Suite [3]), and DRIMust [4]. For example, HOMER could successfully detect the enrichment of the abscisic acid response element (ABRE, ACGTG(G/T)C [5]) in promoters of dehydration-induced genes in pearl millet [6], and the enrichment of the VIP1-binding element (VBE, AGCTG(G/T) [7, 8]) and CAMTA-binding element (CGCGT [9]) in promoters of mechanical stress-responsive genes under the control of the basic leucine zipper (bZIP) transcription factor VIP1 and its close homologs in *Arabidopsis thaliana* [10]. However, those existing tools share four major bottlenecks that hinder a streamlined, comprehensive analysis. First, preparing input data, such as extracting promoter sequences from reference genomes and converting files into tool-specific formats (e.g., FASTA or custom tables), is often technically demanding and requires other tools. Second, many tools are restricted to either command-line interfaces (CLI) on Linux/Unix environments requiring skills or remote web servers, which introduce queuing delays, file size limits, and data privacy concerns when handling unpublished datasets. Third, each tool typically supports only a single analytical paradigm (e.g., hypergeometric (HG) or Fisher’s exact test for HOMER and STREME; minimum HG (mHG) based on a gene expression-ranked list for DRIMust), forcing users to switch between multiple platforms and reformat datasets to evaluate motifs from different perspectives. Last, and most importantly, conventional tools overlook quantitative relationships between motif copy numbers and continuous gene expression measurements: Both HOMER and STREME rely on binary comparisons between target and background sequences; DRIMust uses the expression-ranked list but does not use motif counts.

To overcome these limitations, we developed MEAM-OWL (Motif Enrichment Analysis Modules on the Web, Lite), a lightweight, web browser application. Because MEAM-OWL is a browser application based solely on the native browser language HTML/JavaScript, it has a graphical user interface (GUI) for intuitive data handling and can execute analyses completely within users’ local environments, even offline. For the analysis, it offers five modules, which consist of HOMER-like (Poisson-based approximation of HG), DRIMust-like mHG, Mann– Whitney U (for motif count comparisons between target and background sequences), Extended Core (for Mann–Whitney U-based identification of enriched 4-mer motifs followed by their extension), and Pearson correlation (between motif count and metrics such as gene expression) tests. These analyses can be preceded by sequence extraction and ingestion from genome and genome annotation files (in FASTA and GFF3 formats, respectively), and can be followed by annotation of detected motifs with known *cis*-elements from three public databases, PLACE [11], JASPAR [12] and CIS-BP [13]. The positive correlation between gene induction levels and ABRE counts in promoters in dehydrated pearl millet plants, which had been detected by a custom CLI analysis pipeline [6], was successfully reproduced by the Pearson-based module on MEAM-OWL. Furthermore, the same module detected novel quantitative relationships between gene induction levels and VBE counts in promoters across mechanically stressed Arabidopsis plants where expression of *VIP1* and its close homologs was perturbed. These results demonstrate that the MEAM-OWL integrated platform can promote discoveries of biologically meaningful regulatory motifs and seamlessly connect them to hypothesis building and wet-lab validation.

## 2 Materials and methods

### 2.1 System architecture and implementation

MEAM-OWL is developed solely with HTML5, JavaScript and runs locally in a web browser. The interface, computational routines, genome-reading functions, and multiple-testing adjustment are implemented in index.html. Reference known motifs from PLACE [11], JASPAR [12], and CIS-BP [13] were downloaded and bundled in motif_db.js by a Python script. The pako.min.js was integrated to support decompression of gzip-compressed genome and annotation files. Input processing and statistical calculations are based on five modules classified into two modes, Mode A (target-versus-background) and Mode B (sequence-versus-metrics) (detailed below). For all the modules, *P* values are adjusted using the Benjamini–Hochberg (BH) method [14] and motifs are displayed in ascending order of their *P* values. The overview of the GUI is presented in Fig. 1. The source code is available at GitHub and its archived code is available at the Zenodo repository [15].

**Figure 1.**
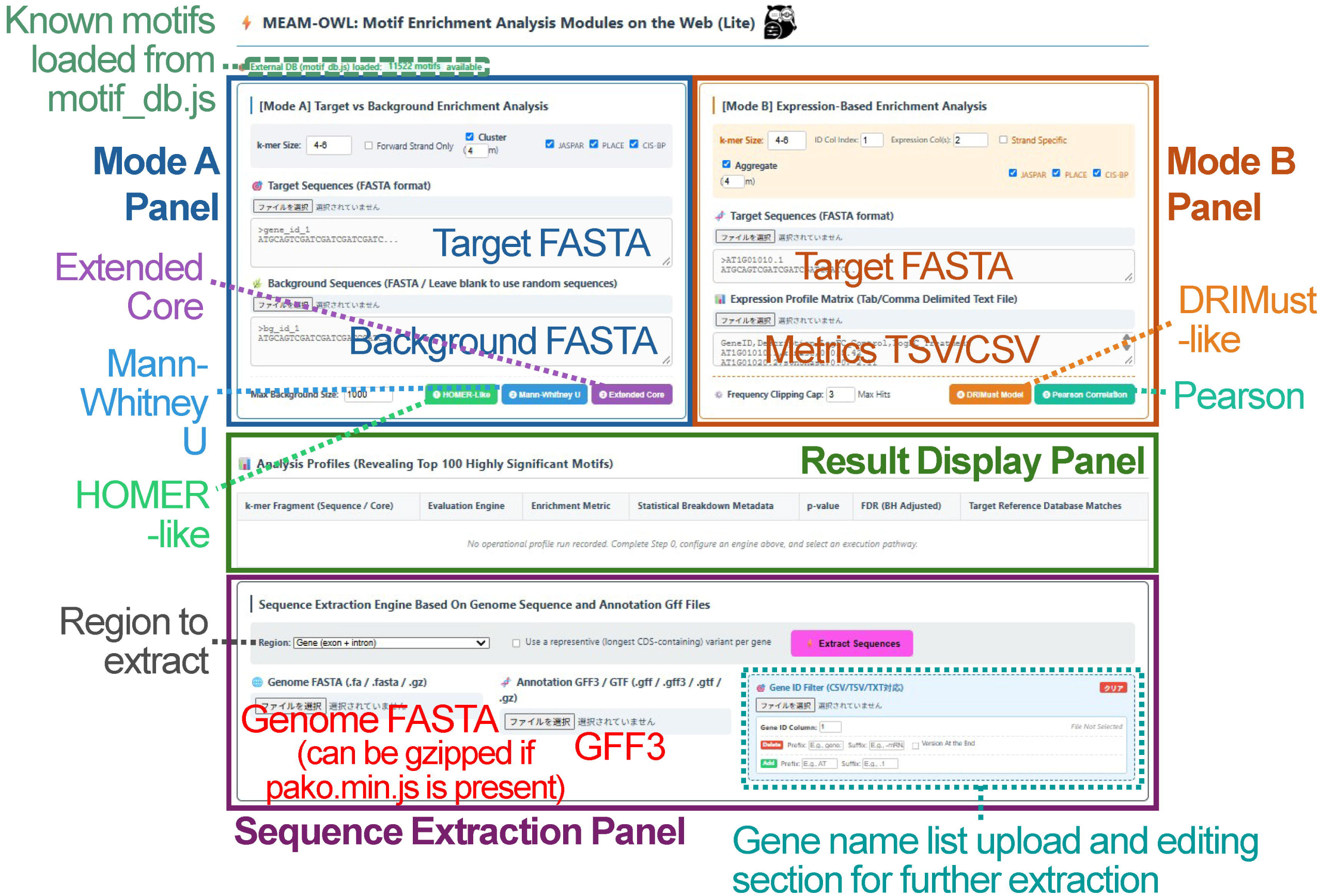
The interface and dependencies of MEAM-OWL. The main analysis functions of MEAM-OWL are written on a single HTML/JavaScript file, allowing it to serve as a single-page application (SPA). Outside files (motif_db.js and pako.min.js) are used to enable comparisons of detected motifs with known motifs and handling gzip-compressed files, respectively.

#### Input data

Users can submit prepared FASTA sequences or extract genomic regions using reference genome (FASTA) and annotation (GFF3) files. Genome files are processed in chunks, retaining only the regions needed for extraction, according to the options provided on the GUI. cDNA sequences are reconstructed from exon annotations. When exon annotations are unavailable, annotated mRNA coordinates can be used as transcript boundaries for extraction. In Custom mode, users can extract regions spanning the specified upstream and downstream distances from the anchor points such as cDNA 5’ ends. Sequences are converted to uppercase and scanned using overlapping windows at selected *k*-mer lengths. Windows containing non-ACGT characters are excluded. Presence/absence and copy number of each *k*-mer are recorded per sequence. Unless strand-specific analysis is selected, occurrences of a *k*-mer and its reverse complement are counted together.

#### Mode A: HOMER-like module

The HOMER-like module compares the fraction of motif-containing sequences between target and background. Enrichment is calculated by dividing the proportion of target sequences containing the motif by the proportion in the background. If a motif is absent in background, the background frequency is set to 1/(N_B_ + 1) (N_B_: the number of background sequences). The expected number of target sequences containing the motif is calculated from background rate. Statistical significance is evaluated with Poisson upper-tail probability, calculated by direct cumulative summation when both the observed and expected counts are ≤20, and by a continuity-corrected normal approximation when either exceeds 20. This module is a simplified implementation rather than a direct execution of the original HOMER algorithm.

#### Mode A: Mann–Whitney U module

The Mann–Whitney U module compares motif counts per sequences between target and background. *P* values are estimated using a one-sided normal approximation with corrections for tied counts and continuity, with tied observations assigned average ranks. Enrichment is the mean target count divided by the mean background count. When the background mean is zero and the target mean is positive, the reported ratio is infinite, displayed as ∞.

#### Mode A: Extended Core module

The Extended Core module uses the same count-based tests in Mann–Whitney U module to identify longer sequences containing short cores. Starting from enriched 4-mer motifs, it evaluates progressively longer *k*-mer containing the current motif and selects the sequence with the smallest *P* value. Extension stops when no eligible candidate is available or the smallest extension *P* value exceeds 0.05. A final candidate is reported only if its BH-adjusted *P* value is below 0.05.

#### Mode B: DRIMust-like module

The DRIMust-like module ranks sequences by their corresponding expression measurements and searches for motif enrichment at both ends of the ranked list. Motifs present in fewer than two sequences or in all sequences are excluded. Interval significance is estimated using a continuity-corrected normal approximation with hypergeometric variance. For *N* matched records, the cutoff step *s* is one when *N* ≤ 1,000; otherwise, it is *N*/200 rounded down to the nearest integer. Each end is searched up to the final cutoff *L* = ⌊0.95*N*⌋, evaluating every *s* position and the endpoint *L*. The minimum *P* value across both searches is multiplied by 2⌊*L/s*⌋ and capped at one. Candidates with a concentration factor above 1.05 and a search-adjusted *P* value below 0.05 proceed to BH adjustment. The concentration factor is the proportion of sequences containing the motif in the selected interval divided by that in the full matched population. A negative sign denotes enrichment at the low-value end.

#### Mode B: Pearson module

The Pearson module pairs motif counts per sequences with metrics (or measurements) such as gene expression levels. Counts exceeding the user-selected cap are replaced with that cap before calculating the Pearson correlation coefficient. Each measurement column, which can be designated through the GUI, is analyzed independently. For *N* valid sequence–measurement pairs, two-sided *P* values are calculated using a Student’s *t* distribution with *N* - 2 degrees of freedom. If all motif counts or all expression measurements are identical, the program sets *r* = 0 and *P* = 1.

#### Result grouping and annotation

Optional grouping organizes retained *k*-mers by shared core sequences. A grouped row inherits the effect value, *P* value, and BH-adjusted *P* value of its member with the smallest *P* value. For grouped results, Mode B distribution plots use the motif with the smallest *P* value in the group. For individual results, plots use the selected motif. Enriched motifs are compared with sequences from the databases using IUPAC consensus compatibility, allowing shorter motifs to match compatible portions of longer ones. Annotations are independent of statistical testing. Results can be exported as CSV, including up to five compatible database records per row in database order rather than confidence order.

#### Multiple-testing adjustment

In Mode A, the testing comprises observed *k*-mers from target and background sequences across the selected lengths, before effect-based filtering. In Mode B, each measurement column is adjusted separately based on the total number of tested *k*- mers across selected lengths. Although the DRIMust-like module applies concentration and cutoff-search *P* value filters before BH adjustment, the full number of *k*-mers is retained for the correction. Final results require a BH-adjusted *P* value below 0.05 and target enrichment in Mode A, a concentration factor greater than 1.05 in DRIMust-like, or absolute Pearson coefficient greater than 0.01. Extended Core also requires the final sequence to pass the BH-adjusted threshold.

### 2.2 Validation of functionality with empirical datasets

To validate the functionality of MEAM-OWL, previously analyzed RNA sequencing (RNA-seq)-derived data for read counts and differentially expressed genes (DEGs) from pearl millet roots exposed to polyethylene glycol (PEG)-induced dehydration for 144 hours (“PRJNA766308-test4”) compared to the unexposed plants, as well as the promoter sequences of these DEGs, were obtained from the Figshare repository [16, 17]. These data were selected because they had led to the detection of the positive correlation between ABRE counts and the gene induction levels as well as the ABRE enrichment in the DEG promoters [6]. RNA-seq data from *Arabidopsis thaliana* leaves of wild-type, QM2 (*VIP1 bZIP29 bZIP30 POSF21* quadruple knockout mutant) and VIP1-GFPox (VIP1-GFP fusion protein-overexpressing) plants were obtained as supplementary data in a previous study [10]. These data were selected because they had led to the detection of the VBE enrichment in the promoters of downregulated genes in QM2 and those of upregulated genes in VIP1ox compared to the wild type [10]. Differential expression analysis (DEA) with these read count data was performed on the DESeq2 [18] engine on DEAR-OWL [19] according to Supplementary Table S1. Resulting lists of DEGs (for both upregulated and downregulated genes) and differential expression levels for all genes were downloaded as CSV files. Pearl millet genome and genome annotation files were downloaded from TGIF-DB [20, 21]. Arabidopsis genome and genome annotation files were downloaded from Ensembl Plants [22]. The DEA-derived files and the genome and genome annotation files were used for the analysis on MEAM-OWL using the HOMER-like, Mann-Whitney U, DRIMust-like and Pearson modules as shown in Fig. 2 and Supplementary Fig. S1. Files with detected motifs were downloaded as CSV files. In the analysis with the Pearson module, violin plots relevant to ABRE and VBE were screenshot to generate Fig. 3. The source and result files necessary for reproducibility were deposited in the Figshare repository [23].

**Figure 2.**
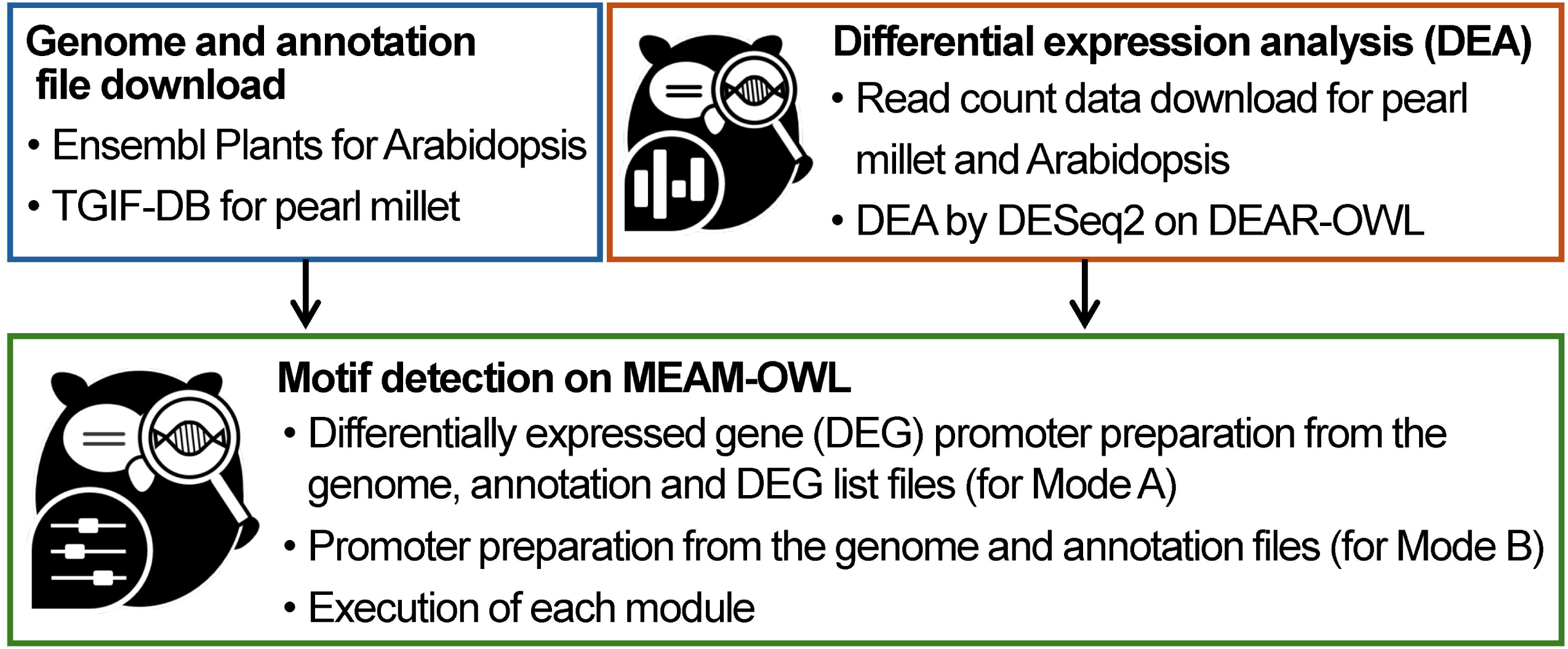
Overview of the validation of MEAM-OWL functionality with empirical datasets. Distribution of log fold change (log_2_FC) for transcripts in the VIP1-GFPox-upregulated DEGs grouped by ACAGCT copy number per promoter. Fold change represents gene expression in mechanically stressed VIP1-GFP-overexpressing plants relative to mechanically stressed wild-type control. X-axis categories indicate motif occurrences per promoter (0, 1, 2, or ≥3), and *n* indicates the number of promoters in each category. Horizontal black bars mark the middle number. Figure shows zoomed y-axis ranges, full-range distributions are provided in Supplementary Figure S2. The ACAGCT distribution was visualized after temporarily removing the top 100 display limit, analysis settings and statistical results were unchanged.

**Figure 3.**
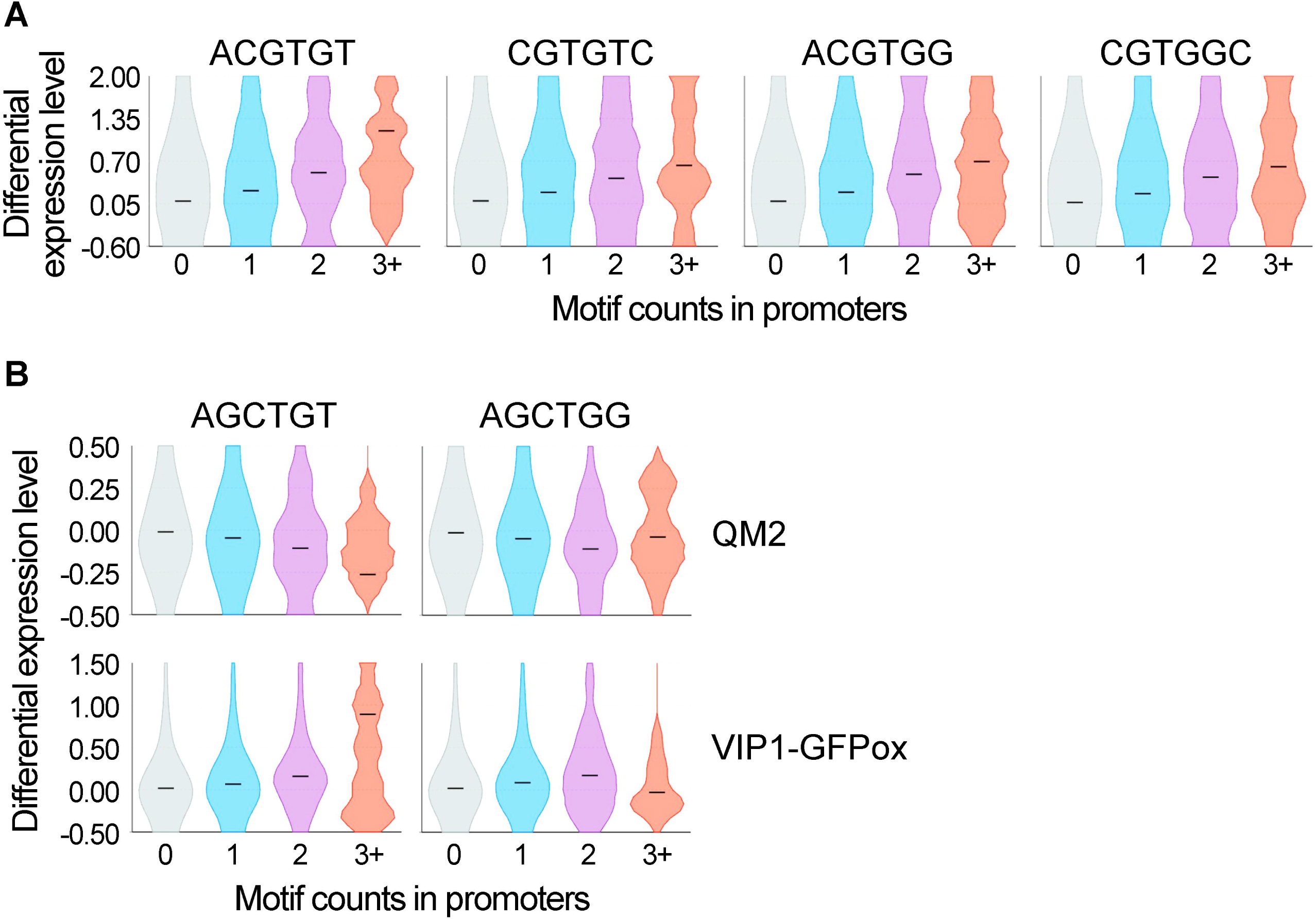
Quantitative correlations between differential gene expression levels and promoter motifs detected by the Pearson module on MEAM-OWL. Violin plots obtained from the Pearson module on MEAM-OWL are presented. The black bars in these plots represent medians. Differential gene expression levels presented in *y*-axes are log fold changes, and “3+” in *x*-axes designates “three or more” in both panels A and B. (A) Positive correlations between differential expression levels and ABRE counts in promoters in dehydrated pearl millet roots. The sequences presented at the top are a part of ABRE (ACGTGGC and ACGTGTC). (B) Negative correlations between differential expression levels and VBE counts in promoters in brushed Arabidopsis QM2 leaves (top panel) and positive correlations between them in brushed VIP1-GFPox leaves (bottom panel).

## 3 Results

### 3.1 Architecture and GUI

All of the five (HOMER-like, Mann–Whitney U, Extended Core, DRIMust-like and Pearson) modules on MEAM-OWL were implemented as relatively short, simple functions using the native browser language HTML/JavaScript. MEAM-OWL can function as a single-page browser application (SPA) with all of those functions included to detect motifs independently of any outside library or file. With the help of outside JavaScript files (pako.min.js and motif_db.js), MEAM-OWL can use gzip-compressed genome (FASTA) and genome annotation (GFF3) files to extract sequences of interest and compare detected motifs with known motifs in databases. Its GUI consists of four panels, Mode A Panel, Mode B Panel, Result Display Panel, and Sequence Extraction Panel, from the top to the bottom (Fig. 1). The Sequence Extraction Panel is optional but allows users to prepare sequences of interest on the fly and to download them.

### 3.2 Validation with empirical datasets

ABRE and VBE had been identified as biologically important motifs in dehydrated pearl millet roots and brushed Arabidopsis leaves with perturbed expression of *VIP1* and its homologs, respectively [6, 9], and were targeted by MEAM-OWL to validate its functionality (Fig. 2).

#### 3.2.1 Pearl millet dataset

In Mode A of MEAM-OWL, using 7-mer motifs as the targets in 500-bp promoters of dehydration-upregulated genes in pearl millet roots, the Extended Core module failed to detect ABRE, but the HOMER-like and Mann-Whitney U modules detected ABRE within the top 10 most significantly enriched motifs (i.e., the motifs with the first to tenth lowest BH-adjusted *P* values) (Supplementary Tables S3 and S4). The fold changes in the occurrences of the two sequences consisting of ABRE (ACGTGGC and ACGTGTC) against the background were 2.32 and 2.35, respectively, in the Mann-Whitney U module (Supplementary Table S4, “Score_index”), consistent with the idea that the ABRE counts in the promoters are important. In Mode B, which limits the maximum motif length to 6-mer for practically short computational time, one partial sequence of ABRE, ACGTGT (or its reverse-complemented sequence ACACGT), was detected within the top 10 most significantly enriched motifs by the DRIMust-like module (Supplementary Table S5), and all the ABRE partial sequences (ACGTGT, CGTGTC, ACGTGG and CGTGGC) were detected by the Pearson module (Supplementary Table S6). The positive correlations between the ABRE counts and differential expression levels in the Pearson module were confirmed by the violin plots (Fig. 3A).

#### 3.2.2 Arabidopsis datasets

In both the Arabidopsis QM2 (where *VIP1* and three of its close homologs were knocked out)-derived and VIP1-GFPox (where VIP1-GFP fusion protein was overexpressed)-derived datasets, while the Extended Core and DRIMust modules failed to detect VBE (either AGCTGG or AGCTGT), all the other modules (i.e., HOMER-like, Mann-Whitney U and Pearson) detected both of the VBE sequences within the top 10 most significantly enriched motifs (Supplementary Tables S3, S4 and S6). In the Mann-Whitney U module, the fold changes in the occurrences of AGCTGG and AGCTGT (or ACAGCT) against the background were 3.68 and 3.20, respectively, for the QM2 dataset, and 2.27 and 1.96 for the VIP1-GFPox dataset (Supplementary Table S4). In the violin plots in the Pearson module, the set of genes with 3+ (three or more) counts of AGCTGT in promoters exhibited the lowest median differential expression level in the QM2 dataset and the highest median differential expression level in the VIP1-GFPox dataset (Fig. 3B, left panel). However, this was not the case for the set of genes with 3+ counts of AGCTGG (Fig. 3B, right). The Pearson module also detected 6-mer motifs containing the CAMTA-binding element CGCGT in both the QM2 and VIP1-GFPox datasets (Supplementary Table S6).

## 4 Discussion

The analysis of the empirical datasets confirms that MEAM-OWL can detect biologically meaningful sequence motifs in provided datasets. The Mann-Whitney and Pearson modules are both specific to this platform and successfully detected the sequences of interest in both the pearl millet and Arabidopsis datasets (Fig. 3 and Supplementary Tables S4 and S6). The DRIMust-like module detected sequences different from ABRE or VBE as the most significant (Supplementary Table S5). These findings demonstrate that different algorithms can detect different motifs from the same dataset provided. The quantitative relationships between the VBE counts and differential expression levels (Fig. 3B and Supplementary Table S6) are new findings obtained by the Pearson module on MEAM-OWL. The analysis with this module also raises the possibility that AGCTGG and AGCTGT differ in contribution to responsiveness to gene expression regulated by VIP1 and its close homologs in vivo (Fig. 3B), unlike in vitro, where those sequences are similarly bound by VIP1 and its close homologs [24, 25]. All of these results demonstrate that integrating multiple algorithms can promote new discoveries and help develop hypothesis for experimental validation. MEAM-OWL serves as such an integrated platform, at the same time securing the data privacy and analysis simplicity and speed as a browser application.

## Acknowledgements

The authors appreciate the computational resources provided by The University of Tokyo.

## Funding

This work was supported by Japan Society for the Promotion of Science (JSPS) KAKENHI Grants (25K09059 and 24KJ0912).

## Declaration on the use of generative AI

During the preparation of this manuscript and the development of the application, the authors utilized a large language model (Gemini 1.5 Flash, Google) to generate the initial structural outlines, drafts of the source code, and documentation, with extensive reviewing and assistance for the backend functions, frontend user interface implementation and writing adjustments. Following this AI-assisted phase, the authors further reviewed, edited, and verified all code components and text to ensure scientific accuracy, technical validity, and compliance with academic standards, taking full responsibility for the final content of the work.

## Software and data availability

The MEAM-OWL application can be executed at https://tsugama-eng.github.io/meam-owl/. Its source code is available on GitHub (https://github.com/tsugama-eng/meam-owl), and its archived release is available on the Zenodo repository [15]. The data used for and obtained from the analysis are listed in Supplementary Table S2, and relevant new data were deposited in the Figshare repository [23].

## Author contributions

D.T., K.K., S.W.A. and X.J. conceived the study and implemented the platform. X.J. wrote the original draft, and J.X., S.W.A., K.K. and D.T. reviewed and edited the manuscript. D.T. and K.K. acquired the funding for this study.

## Competing interests

The authors declare no competing interests.

## Notes

https://doi.org/10.6084/m9.figshare.35008681

### Competing Interest Statement

The authors have declared no competing interest.

