## Supplementary Fig. S1 for "MEAM-OWL for browser-based analysis of motif enrichment and associations with gene expression"

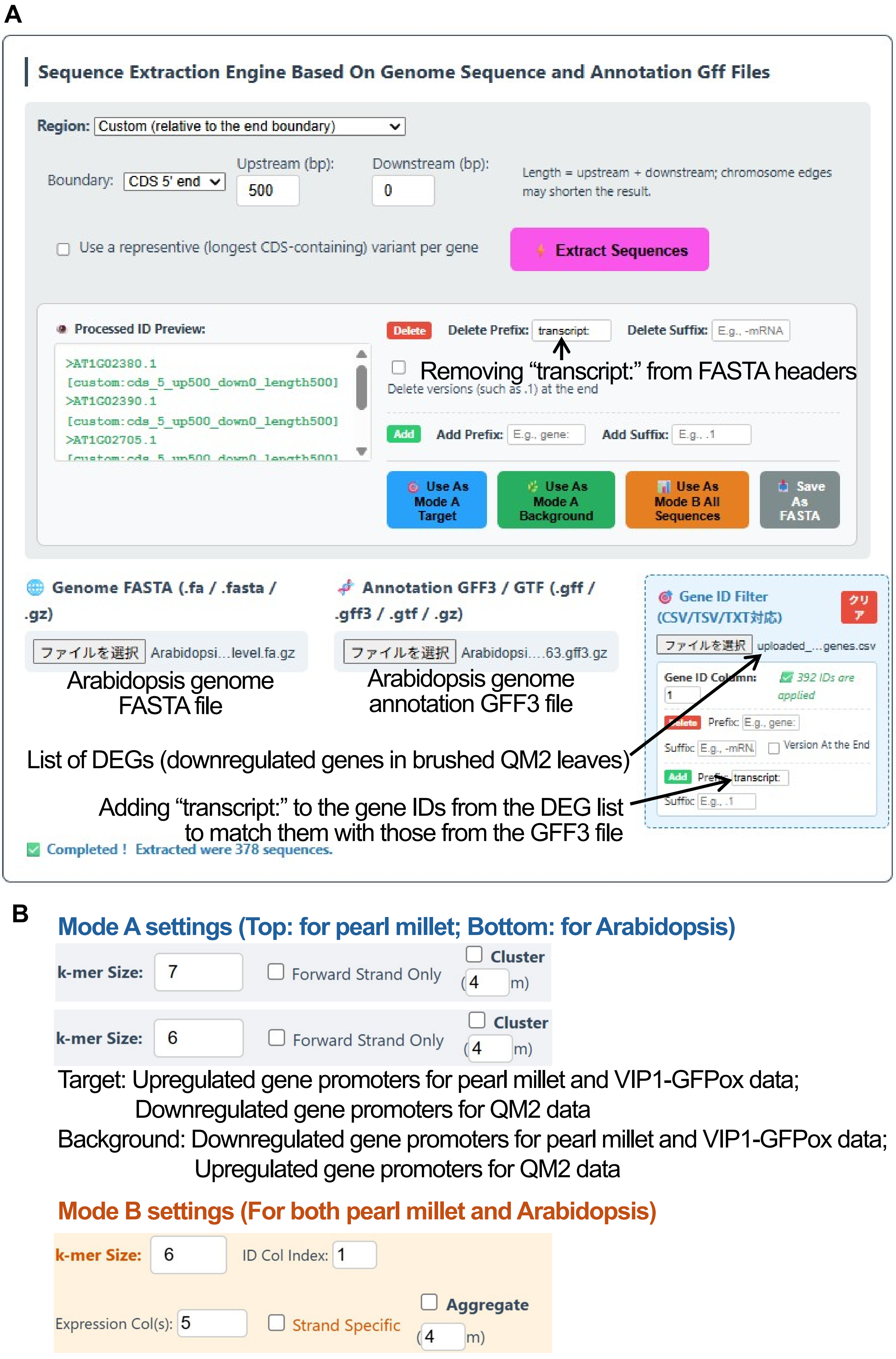


**Supplementary Figure S1. Settings for testing MEAM-OWL for its functionality.** (A) Example settings for preparation of differentially expressed gene (DEG) promoters from genome (FASTA), genome annotation (GFF), and DEG list files. (B) Settings for Mode A (top panel) and Mode B (bottom) for analyses on MEAM-OWL performed in this study.
